# A Python Package for the Localization of Protein Modifications in Mass Spectrometry Data

**DOI:** 10.1101/2022.04.04.487044

**Authors:** Anthony S. Barente, Judit Villén

## Abstract

Determining the correct localization of post-translational modifications (PTMs) on peptides aids in interpreting their effect on protein function. While most algorithms for this task are available as standalone applications or incorporated into software suites, improving their versatility through access from popular scripting languages facilitates experimentation and incorporation into novel workflows. Here we describe pyAscore, an efficient and versatile implementation of the Ascore algorithm in Python for scoring the localization of user defined PTMs in data dependent mass spectrometry. pyAscore can be used from the command line or imported into Python scripts and accepts standard file formats from popular software tools used in bottom-up proteomics. Access to internal objects for scoring and working with modified peptides adds to the toolbox for working with PTMs in Python. pyAscore and is available as an open source package for Python 3.6+ on all major operating systems and can be found at pyascore.readthedocs.io.

## Introduction

Post-translational modifications are fundamental for fine tuning protein function and can be studied at the proteome scale with mass spectrometry (MS). In a typical bottom-up proteomics experiment, whole protein extracts are proteolyzed and the peptides are measured by MS to obtain their mass and fragmentation pattern.^1^ Software tools for protein sequence database search, such as Comet, perform well at matching MS/MS fragmentation spectra to peptide sequences, and identifying when a modification is present in the sequence.^2^ However, they have poor sensitivity at identifying the precise site of modification when the peptide contains multiple acceptor residues.^3^

The Ascore algorithm was one of the first tools to explicitly score the confidence of PTM localization for a peptide-spectra pair.^4^ It is able to provide a probabilistic score for localization confidence by scoring the presence of site-determining ions, i.e. fragment ions that report for the presence of a modification on the specified site. One advantage of this score is that it is calculated and can be consistently interpreted at the level of individual PSMs, and thus can be used in intelligent MS data acquisition workflows to inform subsequent MS events.^5^

The original implementation of Ascore focused solely on phosphorylation and was only available to the community via a web server which restricted submissions to 500 PSMs, a service which itself is now unavailable.^4^ Similarly, a current open source implementation exists within the pyOpenMS software suite, but is limited to phosphopeptides analyzed with CID or HCD type fragmentation and requires the use of suite specific data structures.^6^ There has been growing interest in the proteomics community in analyzing other PTMs and current datasets report 10,000-100,000s spectra assigned to modified peptides. Thus, we reasoned it would benefit the community to expand the capabilities of the Ascore algorithm and make it more broadly accessible, so that it can be used as an alternative to or in conjunction with other open source PTM localization algorithms.^7^ Here we present pyAscore, a fast and extensible open source implementation of the Ascore algorithm, which can handle a wide variety of modifications and MS/MS fragmentation modes.

## Methods

Mass spectrometry data for individual experiments was downloaded directly from MassIVE or PRIDE, converted to mzML format with ThermoRawFileParser (v. 1.3.4) and then searched with the Comet database search software (v. 2021010).^2^ Recommended parameters for high resolution and low resolution searches were taken from the Comet documentation. Human samples were searched with the uniprot *Homo sapiens* reference proteome (downloaded Feb 8, 2022), and all synthetic peptide data was searched with the FASTA from PXD000138. All files were searched with carbamidomethylation on cysteines as static modification and oxidized methionine as variable modification. Since dataset PXD007145 reports TMT-labeled peptides, a TMT 10-plex modification was added as a static modification on lysines and peptide N-termini. For the synthetic peptide datasets, PXD000138 and PXD000759, and the other phosphoproteomic datasets, PXD007740 and PXD007145, the modification of interest was phosphorylation on serine, threonine, and tyrosine, and this was included in the variable modification list. For dataset MSV000079068, acetylation of the protein N-terminus and internal lysines was included as a variable modification, but peptide C-terminal lysines were not allowed to be acetylated. Finally, HCD and CID data were searched with the b and y ion series, whereas ETD data was searched with the c and z+H ion series. All searches were subsequently grouped by dataset and fragmentation method and analyzed with Mokapot using default parameters.^8^ Unless otherwise noted, for pyAscore localization, the fragment error was set to 0.05 Da and the same ion series as the searches was used. All searches and tests were performed on Intel Xeon Gold 6312U 2.4Ghz processors.

## Results and Discussion

### Overview of package and implementation

pyAscore provides an accessible interface to perform probabilistic scoring of PTM localization directly from the command line or incorporated into Python scripts (Fig. 1a). In both cases, our package relies on pyteomics to provide the ability to read from popular file formats containing mass spectra and peptide spectrum matches (PSMs),^9^ so that scoring can be incorporated into a wide variety of workflows. For each input of MS/MS spectrum, peptide sequence, and a modification of interest, pyAscore will provide the overall best localization and an ambiguity score, which describes how much better the best localization is compared to the next best alternative. This procedure generalizes to any modification of interest just by specifying the mass of the modification and the potential acceptor amino acids, e.g. 79.9663 Da and Ser, Thr, Tyr in the case of phosphorylation. Users can also customize analytical parameters to their specific MS acquisition method, with parameters for fragment ion mass tolerance, ion types to score, and presence of neutral losses.

**Figure 1.**
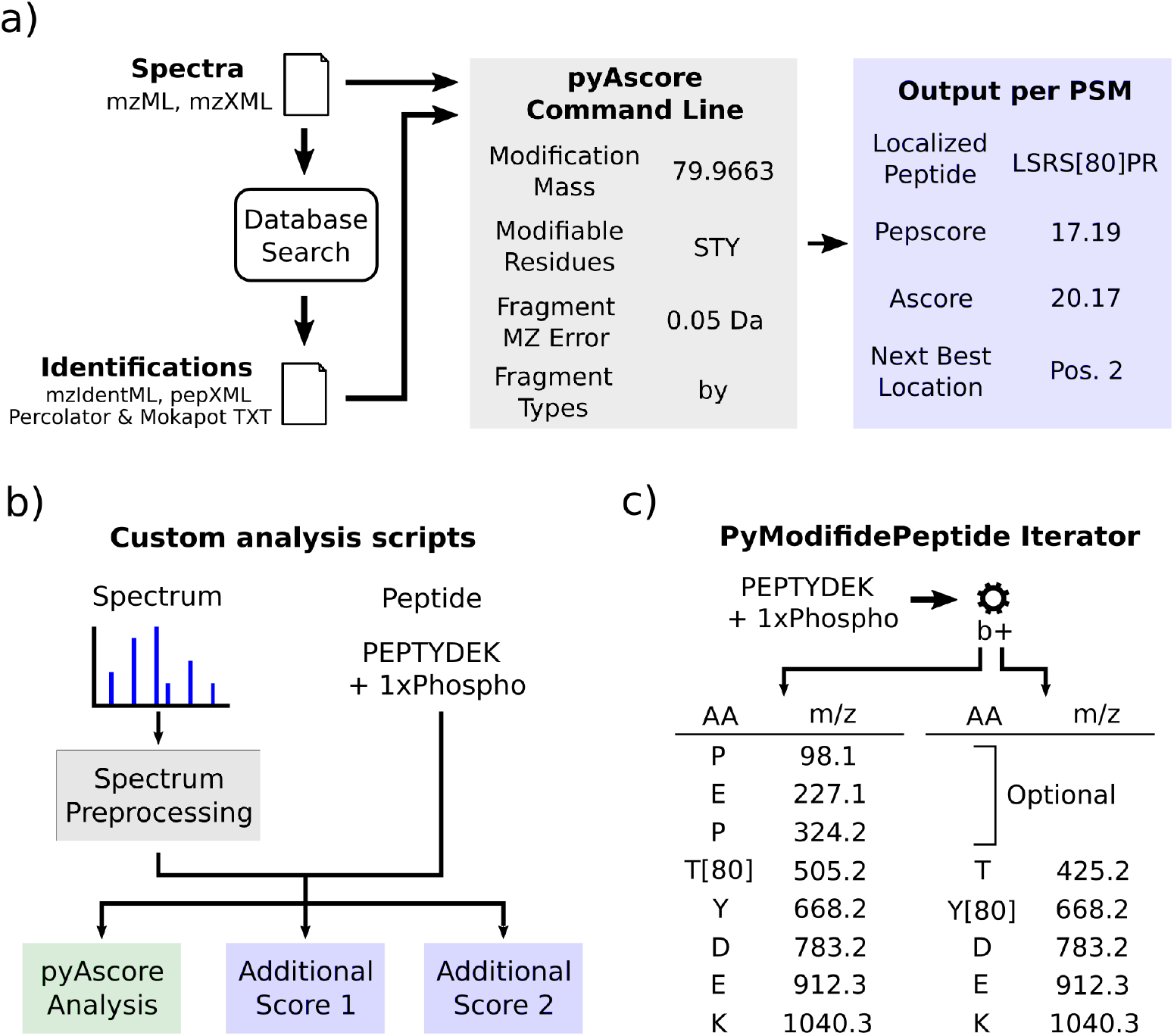
**a)** pyAscore can be used directly from the command line as a standalone application with a user specified modification and instrument parameters. The required input is a file containing spectra and a file containing peptide spectrum matches, both of which can be supplied via multiple popular mass spectrometry data formats. pyAscore outputs localization information per modified PSM in a tsv file. **b)** Individual scans and PSMs can be passed to pyAscore in Python scripts, allowing on the fly logic and the combination of multiple analyses per PSM. **c)** pyAscore provides Python use of its internal iterator pyModifiedPeptide, which allows users to efficiently step through theoretical fragment masses for all permutations of a peptide and a modification, with the b series ions shown here.

The core algorithmic components of pyAscore are implemented in C++ and exposed to Python using the Cython package. This facilitates the production of fast and efficient code for scoring while allowing the flexibility of analyses provided by the Python programming language. The full scoring procedure and score definitions correspond to the original descriptions by Beausoleil et al.,^4^ and we briefly describe the main scoring steps here. First, the MS/MS spectrum is binned into 100 m/z windows and the fragment ions within each bin are ranked. pyAscore then iterates over every possible localization of the PTM of interest on the peptide backbone and produces a PepScore, which is based on the number of matching theoretical peaks between a modified peptide sequence and the ranked set of fragment ions. The best scoring localization is then reported to the user. Peptides can potentially have a large number of permutations of unmodified and modified sites, so care was taken at this step to increase speed by reducing repeat fragment mass calculations and corresponding peak search.

Finally, for each modified site on the best localized peptide sequence, pyAscore calculates a score based on the number of site-determining ions for the best localization according to the Pepscore *vs*. the next best localization with that site unmodified. Like in the original development^4^, this score is termed the ambiguity score, or Ascore, and gives a probabilistic metric at how much better the best localization is than the next best possible modification site.

If more intricate analyses are desired, the components of pyAscore can be imported into Python scripts or any software that can link C++ libraries. Users then have direct access to the internal scoring class, which can be combined with spectrum preprocessing or further localization scoring (Fig. 1b). Since multiple scoring objects can be created with their own specific parameters, users also have the option to tailor score calculation to individual scan types. For users in need of tools for fast prototyping of new localization algorithms, pyAscore also provides access to its internal class for iterating over permutations of modified residues on peptides and calculating their individual theoretical fragment masses (Fig. 1c).

### Speed of localization scoring

To evaluate pyAscore run times and usability in production settings, we reanalyzed three replicate high-resolution data-dependent acquisition (DDA) label-free phosphoproteomic runs from PXD007740.^10^ After database search, pyAscore phosphosite localization was timed for each PSM.^8^ Median localization times for individual replicates ranged from 0.061 ms to 0.073 ms per PSM with 99.7% of localizations taking 1 ms or less (Fig. 2a) and all PSMs in the run being scored in a matter of seconds. Plotting the localization time by the number of localization permutations in logarithmic scale revealed a linear correspondence. This is likely driven by the PepScore calculation, which must perform a calculation for each permutation of phosphorylated and not phosphorylated sites. Despite this, we found that the rate limiting step in the pyAscore calculations for a single file tended to be reading spectra and PSMs into memory.

**Figure 2.**
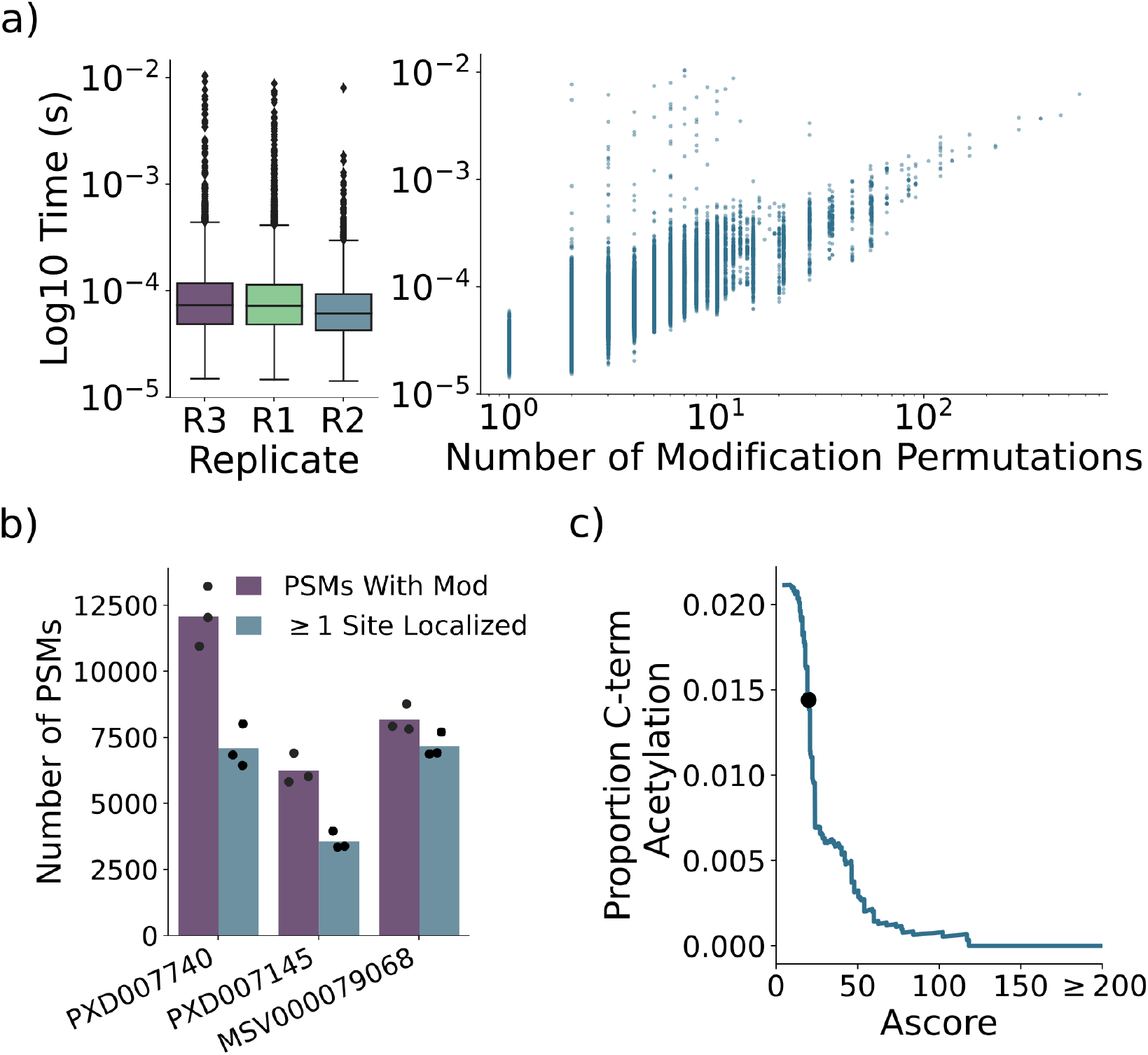
**a)** Analysis time per PSM for 3 replicate phosphoproteomic runs from PXD07740 (left), and the analysis time split by the number of unmodified/modified site permutations (right). **b)** Number of PSMs with a modification and the number of PSMs with at least one site localized for 3 replicate runs from each of PXD007740 (label-free phosphoproteome), PXD007145 (TMT-labeled phosphoproteome), and MSV000079068 (label-free acetylome). **c)** Cumulative number of falsely localized C-terminal acetylations by Ascore for all PSMs containing both an internal lysine and C-terminal lysine from MSV000079068.

**Figure 3.**
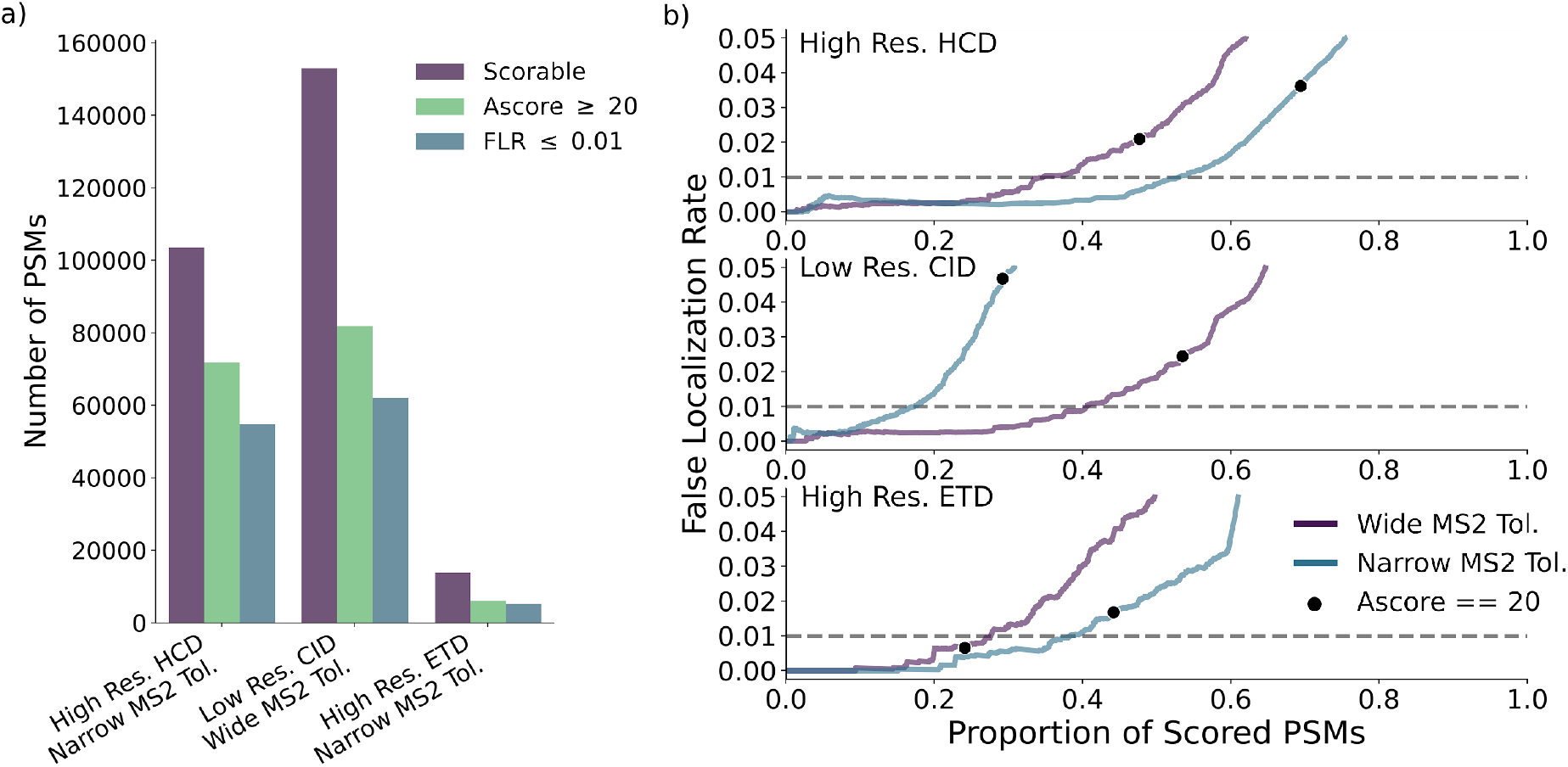
**a)** The number of PSMs which can be scored by pyAscore, the number of PSMs passing an Ascore cut off of 20, and the number of PSMs at a true FLR cutoff of 1% for the Marx *et al*. synthetic peptide dataset measured with 3 fragmentation methods. PSMs were considered scorable if the peptide came from the synthetic library, had a single phosphorylation event, and had at least 2 common phosphorylatable amino acids (i.e. Ser, Thr, Tyr). For each dataset, only the pyAscore results with parameter settings best matched to the instrument acquisition settings are shown, i.e. narrow tolerance settings for high resolution data and wide tolerance settings for low resolution data. **b)** False localization rate vs proportion of scored PSMs at decreasing Ascore cutoffs. Results are shown for pyAscore run with wide fragment mass tolerance (i.e. ±0.5 Da) and pyAscore run with narrow fragment mass tolerance (±0.05 Da). False localization rate (FLR) is calculated as the number of PSMs with Ascore passing at a given threshold where pyAscore called the wrong localization, divided by the total number of passing PSMs.

### Versatility of the pyAscore package

We wanted to showcase the versatility of pyAscore to extend to other modifications. Thus in addition to the data from PXD007740 we analyzed three replicate high-resolution DDA runs from both PXD007145,^11^ which contained TMT labeled phosphopeptides, and MSV000079068,^12^ which contain label-free acetylated peptides. For both PXD007740 and PXD007145, pyAscore was set to localize phosphorylations to serine, theonine, or tyrosine residues, and for MSV000079068 pyAscore was set to localize acetylations to any lysine in the peptide. An Ascore cutoff of 20 (theoretical confidence of 99%) has often been used in previous studies for determining localization, and thus we considered any site which achieved this score as localized. For the label free phosphoproteomics data, pyAscore determined an average of 58.8% of modified PSMs contained at least one localized site (Fig. 2b). The TMT-labeled phosphoproteomic data performed similarly, with an average of 57.1% of modified PSMs containing at least one localized site. On the acetylation dataset, pyAscore determined that an average of 87.7% of modified PSMs contain at least one localized site. This is likely due to the overall lower number of acceptor sites per peptide in the acetylation data than the phosphorylation data.

Our pyAscore analysis of the acetylation data (MSV000079068) was slightly less stringent than our database search, the latter of which did not allow acetylation on C-terminal lysines. This provided a first look into the underlying false localization rate of pyAscore, as we could count any acetylations that pyAscore placed on the C-terminal lysines as incorrect. We thus filtered the PSMs of MSV000079068 for those that contained a single acetylation event, a C-terminal lysine, and at least one other internal lysine (N=14,608). We then sorted the PSMs by decreasing Ascore, and counted the cumulative number of C-terminal acetylations as we moved down the list. This showed a direct correspondence between the Ascore and the rate of C-terminal acetylation, with the Ascore cutoff of 20 achieving a rate of 1.44% C-terminal acetylation (Fig. 2c). While this provides a great deal of confidence that pyAscore is performing well, the fact that these spectra were provided by Comet as ones that already had low potential for C-terminal acetylation led us to look for another independent dataset to directly evaluated pyAscore’s false localization rate.

### Validation of pyAscore on a dataset of synthetic phosphopeptides

We wanted to evaluate the performance of pyAscore’s PTM localizations on data where the modification site of peptides was known. Thus, we turned to a large synthetic phosphopeptide library produced by Marx *et al*. (2013).^13^ This library was analyzed across 2 studies with 3 MS acquisition approaches: high resolution HCD, high resolution ETD, and low resolution CID.^13,14^ The analysis of the same data with different fragmentation modes and resolution settings allowed us to compare how parameter choices in pyAscore affected performance, so we downloaded files from both repositories corresponding to the analysis of the original library.

After database search, we filtered for singly phosphorylated PSMs which were part of the synthetic library and obtained a total of 103,561, 153,088, and 13,854 PSMs in the high resolution HCD, low resolution CID, and high resolution ETD datasets respectively (Fig. 2a). While the number of PSMs was much lower in the ETD dataset than the others, there was still an ample number of PSMs to evaluate FLR. For PTM localization, we decided to evaluate the effect of tailoring pyAscore’s parameters to instrument parameters during scores. Thus, for each dataset we tested two parameter settings for pyAscore’s fragment ion mass tolerance. The wide tolerance, ±0.5 Da, corresponds to the tolerance used in the original Ascore paper, but is presumably too tolerant for high resolution data.^4^ Therefore, we also tested a narrow tolerance setting, ±0.05 Da.

After scoring, we ordered PSMs by decreasing localization score, and evaluated the relationship between the score and the true FLR. For the two high resolution datasets, using the narrow mass tolerance drastically increases the number of PSMs at a given FLR, while the low resolution dataset performed better with the wide mass tolerance (Fig. 2b). This directly shows the benefit of tailoring the scoring parameters to acquisition parameters. It is notable that an Ascore cut off of 20 varied in FLR depending on dataset and parameter settings. When localization parameters did not match acquisition parameters, the FLR at an Ascore of 20 was 2.1%, 4.7%, and 0.7% for the HCD, CID, and the ETD data respectively, while the FLR for tailored parameters was 3.6%, 2.2%, and 1.7% for the same datasets. However, taken together with the acetylation results presented above, these data suggest that an Ascore cutoff of at least 20 can still achieve a consistently low FLR if parameters are tailored correctly. In total, with tailored parameters, pyAscore scored 52.96%, 40.53%, and 38.28% of PSMs at a 1% FLR for the HCD, CID, and ETD respectively, and 69.38%, 53.45%, and 44.15% of PSMs at an Ascore cutoff of 20 for the same data (Fig. 2a).

## Conclusions

Here we describe pyAscore, a flexible open source Python package for probabilistic scoring of PTM localization. Our package can be incorporated into a variety of analytical workflows and provides full access to internal parameters to tailor analyses to the user’s instrument and PTM of interest. We showed that pyAscore is fast enough to be used in high performance settings such as in intelligent mass spectrometry data acquisition alongside real time database search. In our online documentation, we supply examples of potential workflows in order to provide users with a detailed look at how they can apply pyAscore to their own data.

## Data availability

Analyses within this study were performed on publicly available data from the ProteomeXchange^15^ partner repositories, PXD007740, PXD007145, PXD000138, and PXD000759 from PRIDE (https://www.ebi.ac.uk/pride/archive) and MSV000079068 from MassIVE (https://massive.ucsd.edu). Result files from search and scoring steps can be found at the PRIDE repository PXD032908, and all code for reproducing validation experiments and generating figures can be found on GitHub at https://github.com/AnthonyOfSeattle/pyAscoreValidation.

## Acknowledgements

The authors would like to thank Ian Smith and Bianca Ruiz from the Villén lab, and Ralf Gabriels from Lennart Martens’ lab for early testing of pyAscore and providing valuable feedback on features and bugs. The authors would also like to thank members of the Villén lab for their helpful comments and guidance during development and validation.

## References

1. Zhao, Y. & Jensen, O. N. Modification-specific proteomics: strategies for characterization of post-translational modifications using enrichment techniques. Proteomics 9, 4632–4641 (2009).

2. Eng, J. K., Jahan, T. A. & Hoopmann, M. R. Comet: an open-source MS/MS sequence database search tool. Proteomics 13, 22–24 (2013).

3. Chalkley, R. J. & Clauser, K. R. Modification site localization scoring: strategies and performance. Mol. Cell. Proteomics 11, 3–14 (2012).

4. Beausoleil, S. A., Villén, J., Gerber, S. A., Rush, J. & Gygi, S. P. A probability-based approach for high-throughput protein phosphorylation analysis and site localization. Nat. Biotechnol. 24, 1285–1292 (2006).

5. Schweppe, D. K. et al. Full-Featured, Real-Time Database Searching Platform Enables Fast and Accurate Multiplexed Quantitative Proteomics. J. Proteome Res. 19, 2026–2034 (2020).

6. Röst, H. L., Schmitt, U., Aebersold, R. & Malmström, L. pyOpenMS: a Python-based interface to the OpenMS mass-spectrometry algorithm library. Proteomics 14, 74–77 (2014).

7. Shteynberg, D. D. et al. PTMProphet: Fast and Accurate Mass Modification Localization for the Trans-Proteomic Pipeline. J. Proteome Res. 18, 4262–4272 (2019).

8. Fondrie, W. E. & Noble, W. S. mokapot: Fast and Flexible Semisupervised Learning for Peptide Detection. J. Proteome Res. 20, 1966–1971 (2021).

9. Goloborodko, A. A., Levitsky, L. I., Ivanov, M. V. & Gorshkov, M. V. Pyteomics--a Python framework for exploratory data analysis and rapid software prototyping in proteomics. J. Am. Soc. Mass Spectrom. 24, 301–304 (2013).

10. Ressa, A. et al. A System-wide Approach to Monitor Responses to Synergistic BRAF and EGFR Inhibition in Colorectal Cancer Cells. Mol. Cell. Proteomics 17, 1892–1908 (2018).

11. Hogrebe, A. et al. Benchmarking common quantification strategies for large-scale phosphoproteomics. Nat. Commun. 9, 1045 (2018).

12. Svinkina, T. et al. Deep, Quantitative Coverage of the Lysine Acetylome Using Novel Anti-acetyl-lysine Antibodies and an Optimized Proteomic Workflow. Mol. Cell. Proteomics 14, 2429–2440 (2015).

13. Marx, H. et al. A large synthetic peptide and phosphopeptide reference library for mass spectrometry–based proteomics. Nature Biotechnology vol. 31 557–564 (2013).

14. Matheron, L., van den Toorn, H., Heck, A. J. R. & Mohammed, S. Characterization of biases in phosphopeptide enrichment by Ti(4+)-immobilized metal affinity chromatography and TiO2 using a massive synthetic library and human cell digests. Anal. Chem. 86, 8312–8320 (2014).

15. Vizcaíno, J. A. et al. ProteomeXchange provides globally coordinated proteomics data submission and dissemination. Nat. Biotechnol. 32, 223–226 (2014).

